# Multi-Scale Label-free Human Brain Imaging with Integrated Serial Sectioning Polarization Sensitive Optical Coherence Tomography and Two-Photon Microscopy

**DOI:** 10.1101/2023.05.22.541785

**Authors:** Shuaibin Chang, Jiarui Yang, Anna Novoseltseva, Xinlei Fu, Chenglin Li, Shih-Chi Chen, Jean C. Augustinack, Caroline Magnain, Bruce Fischl, Ann C. Mckee, David A. Boas, Ichun Anderson Chen, Hui Wang

**Affiliations:** Department of Electrical and Computer Engineering, Boston University, 8 St Mary’s St, Boston 02215, USA; Department of Biomedical Engineering, Boston University, 44 Cummington Mall, Boston 02215, USA; The Chinese University of Hong Kong, Department of Mechanical Engineering, Hong Kong Special Administrative Region, China; Department of Radiology, Massachusetts General Hospital, A.A. Martinos Center for Biomedical Imaging, 13th Street, Boston 02129, USA; VA Boston Healthcare System, U.S. Department of Veteran Affairs; Boston University Chobanian and Avedisian School of Medicine, Boston University Alzheimer’s Disease Research Center and CTE Center; Department of Neurology, Boston University Chobanian and Avedisian School of Medicine; Department of Pathology and Laboratory Medicine, Boston University Chobanian and Avedisian School of Medicine; VA Bedford Healthcare System, U.S. Department of Veteran Affairs, Bedford, MA, USA

## Abstract

The study of neurodegenerative processes in the human brain requires a comprehensive understanding of cytoarchitectonic, myeloarchitectonic, and vascular structures. Recent computational advances have enabled volumetric reconstruction of the human brain using thousands of stained slices, however, tissue distortions and loss resulting from standard histological processing have hindered deformation-free reconstruction of the human brain. The development of a multi-scale and volumetric human brain imaging technique that can measure intact brain structure would be a major technical advance. Here, we describe the development of integrated serial sectioning Polarization Sensitive Optical Coherence Tomography (PSOCT) and Two Photon Microscopy (2PM) to provide label-free multi-contrast imaging, including scattering, birefringence and autofluorescence of human brain tissue. We demonstrate that high-throughput reconstruction of 4×4×2cm^3^ sample blocks and simple registration of PSOCT and 2PM images enable comprehensive analysis of myelin content, vascular structure, and cellular information. We show that 2 *μm* in-plane resolution 2PM images provide microscopic validation and enrichment of the cellular information provided by the PSOCT optical property maps on the same sample, revealing the sophisticated capillary networks and lipofuscin filled cell bodies across the cortical layers. Our method is applicable to the study of a variety of pathological processes, including demyelination, cell loss, and microvascular changes in neurodegenerative diseases such as Alzheimer’s disease (AD) and Chronic Traumatic Encephalopathy (CTE).

## 1. Introduction

The human brain is organized at multiple scales, from the macroscopic white-grey matter structure to the microscopic components of neurons, axons, vascular structures. Mapping brain structures at different scales is essential for functional and pathological studies. Magnetic resonance imaging (MRI) is widely used for mapping the brain’s macroscopic structure^1–3^, however, it is limited in its ability to achieve the microscopic spatial resolution. Label-free, automatic serial sectioning optical coherence tomography (OCT) is complementary to traditional histology as it generates depth-resolved 3D images and allows intact tissue structure to be reconstructed. It also utilizes intrinsic scattering contrasts to measure tissue structure and composition and has demonstrated utility in mouse and human brain^4–6^. Polarization sensitive OCT (PSOCT) measures the birefringence properties of myelinated fiber tracts to provide additional contrast^7–9^. Wang et al^10^ demonstrated a fully automated pipeline using a serial sectioning PSOCT to image tens of cubic centimeters of brain tissue, enabling quantitative estimation of cortical layer thickness and generation of 2D microscopic images that were useful for orientation in tractography. While PSOCT lacks the resolution and sensitivity to visualize individual neurons, axons, and capillaries, optical coherence microscopy (OCM) is capable of visualizing neurons in human^11^ and mouse brain^12^. Drawbacks of OCT/OCM include the requirement for depth scanning, which slows imaging of large tissue volumes, and speckle noise, that interferes with accurate neuronal counts^11,12^.

Two photon microscopy (2PM) is an optical imaging technique that suppresses out-of-focus fluorescence and generate high-resolution images with excellent signal to background ratio. This technique has been applied to a variety of tissue types, including skin^13,14^, retinal^15,16^, cancerous tissue^17,18^ and brain^19,20^. Multiple endogenous fluorophores are two-photon excitable, such as collagen and elastin fibers^21^, iron-free variants of hemoglobin and macromolecules in blood plasma^22^, and lipofuscin^23^ in aging neurons. The presence of these endogenous fluorophores in postmortem human brain offers opportunities for label-free, high-resolution imaging of neurons, capillaries, and axonal fibers, which can complement the images provided by PSOCT.

Here, we present our efforts to combine serial sectioning PSOCT and 2PM to image large volume human brain samples at multiple scales. Our results show that by integrating the scattering, birefringence and autofluorescence contrasts, we were able to investigate multiple perspectives of vascular structure, myelin content, and neurons. Previously, hybrid systems combining OCT and 2PM were applied in wound healing^24^, microvascular-flow distribution^25^, skin inflammation^14^ and skin diseases^18^. To our knowledge, no previous work has combined serial sectioning, PSOCT, and 2PM in imaging human brain tissue. Our system is capable of simultaneously imaging and registering key features in human brain, opening a window for investigating their spatial interactions and pathological relevance in brain aging and neurodegeneration.

## 2. Results

### 2.1 Serial Sectioning PSOCT-2PM microscope

The serial sectioning PSOCT-2PM system setup is shown in Figure 1(a) and a detailed description is included in the Methods section. The excitation wavelength of 2PM was 820nm and two detection channels were used: one covering 460±25nm (short wavelength channel) for detecting autofluorescence from elastin, collagen, blood, and fixation agent, and the other covering 600±100nm (long wavelength channel) for detecting autofluorescence from lipofuscin. The Field of View (FOV) was 3×3mm^2^ and the lateral and axial resolution were 2 *μm* and 48 *μm* (shown in Supplementary Figure 1), respectively. The PSOCT setup is shown on the right part of Figure 1(a). It has the same 3×3mm^2^ FOV as 2PM, with 5 *μm* isotropic resolution and 150 *μm* depth of focus (shown in Supplementary Figure 1&2). Under the objective, the sample was embedded in 4% agarose and mounted on XYZ motorized stages using a 3D printed base plate (shown in inset) to secure it for measurements spanning multiple days. The stages translated the sample under the objective to cover the whole area, as well as between the vibratome and the objective. To obtain large volumetric imaging, a customized vibratome^26^ equipped with a 2.5inch wide sapphire blade was integrated into the imaging system. The vibratome was designed to minimize out-of-plane vibration for optimal cutting performance^27,28^. The cutting thickness was 150 and 450, respectively, for non-refractive-index-matched samples and refractive-index-matched samples. With our optimized imaging and cutting scheme and data storage, a 4×4×2cm^3^ brain block can be imaged within 6 days.

**Figure 1.**
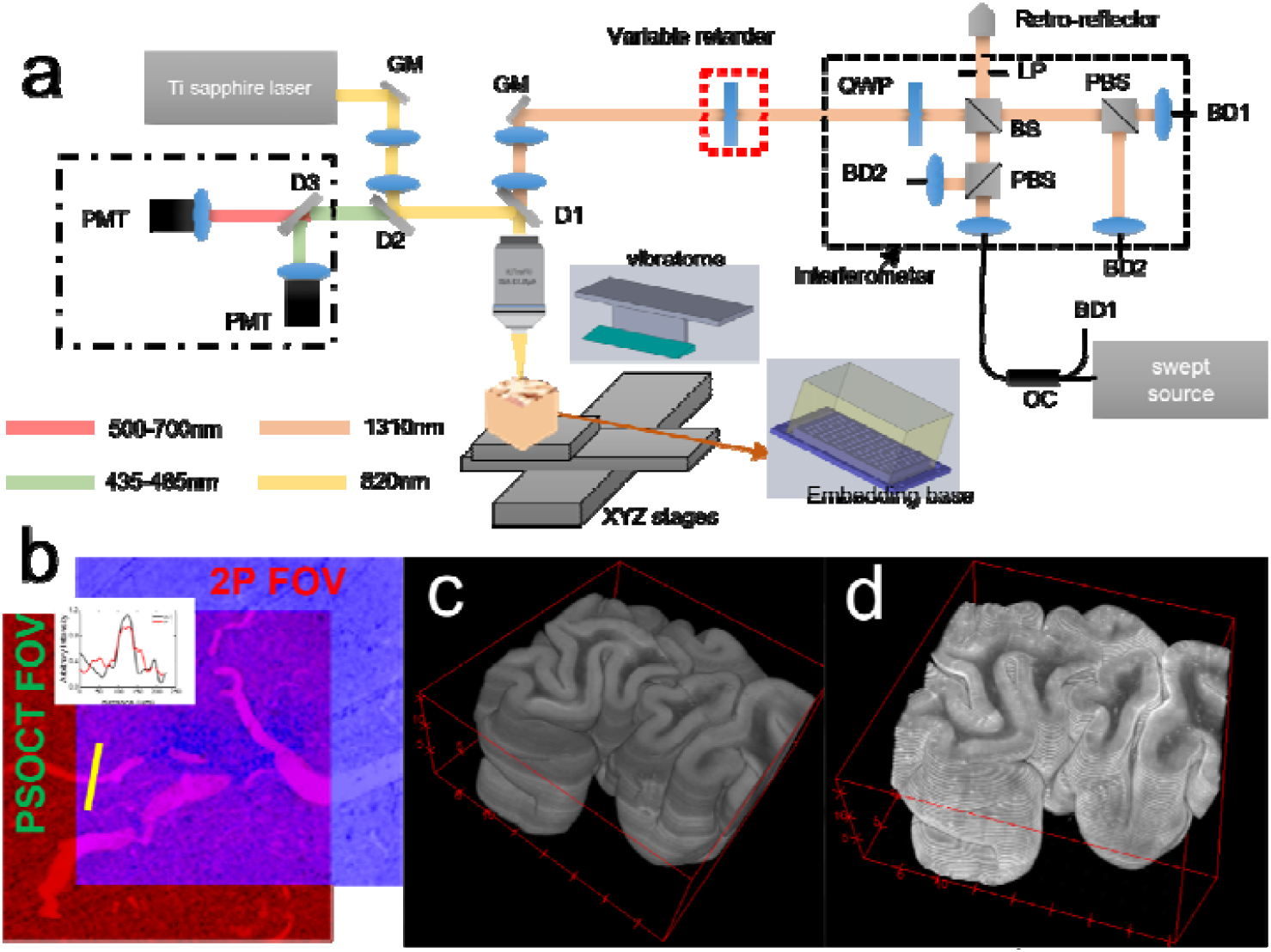
PSOCT-2PM system schematic and volumetric reconstructions from a 4×4×2cm^3^ human brain sample. (a) Optical layout of the hybrid system. PMT: photomultiplier tube. D1-D3: Dichroic mirrors. GM: Galvo mirror pair. QWP: quarter wave plate. LP: linear polarizer. PBS: polarized beam splitter. BS: beam splitter. BD: balanced detectors. OC: optical circulator. The sample embedding is shown in the inset. A 3D printed sample plate was used to hold the whole block and screwed to the bath. (b) Illustration of the overlapping 2PM and PSOCT FOVs. Inset shows the line profile of a vessel in the overlapping area. (c-d) Volumetric rendering of PSOCT (c) and 2PM (d) images of a human brain sample. Both volumes were down-sampled to an isotropic voxel size of 60*μ*m for easier rendering, XYZ axis units were in mm.

We carefully aligned the FOV of PSOCT and 2PM to ensure the maximum overlap and minimized the need for post-processing registration. Since the two modalities are imaged simultaneously, linear translation of the FOVs was sufficient for registration. Figure 1.b show the overlap of the two FOVs for a region with multiple vessels. PSOCT is in red and 2PM is in blue. The co-aligned vessels in the overlap area and their line profiles in the inset show the successful registration between the two modalities. Figure 1.c-d show the volumetric reconstruction of the PSOCT and 2PM images of a 4×4×2cm^3^ sample from Broca’s area 44/45 (BA44/45). The sample was index matched to reduce the scattering and increase the penetration depth of both PSOCT and 2PM. In PSOCT volume (c) white matter appeared as darker than the grey matter due to a higher scattering coefficient. 2PM volume (d) that were registered to PSOCT also presented a contrast between grey and white matter. The autofluorescence signals were stronger in grey matter than white matter, which may be due to the higher concentration of autofluorescent components such as elastin-rich Extra Cellular Matrix^29^ (ECM) and vessels, a well as lipofuscin-filled neurons^23^. Volumetric fly-through of PSOCT and 2PM images are included in supplementary videos.

Birefringence of brain tissue is an important feature exploited by PSOCT to provide contrast that is sensitive to myelin content and axonal orientation. However, when combining PSOCT with 2PM, the optics added to the PSOCT light path such as the dichroic mirror and the telescope introduced a system birefringence, preventing an accurate measure of sample birefringence. To compensate this system birefringence, we used an electrically modulated variable retarder (Thorlabs Inc, red dashed box in Figure 1(a)) that was oriented with the same optic axis but opposite retardance as the optics in the sample arm. Figure 2 shows a comparison of the polarization extinction ratio (PER), dynamic range of retardance and optic axis orientation before and after system birefringence compensation. Before compensation, the PER (Figure 2.a) of co-polarization channels was below 20dB off the center of the FOV, and the PER of cross-polarization channels was below 5dB. After compensation, PER increased by 10-30dB, recovering PSOCT’s ability to measure sample birefringence. Figure 2.b shows the retardance measurement before and after compensation. Before compensation, the hybrid system produced a background retardance of approximately 70° and the measurement on the sample was inaccurate. After compensation, the background was removed and the dynamic range of retardance measurement was recovered to about 90°. The measurement matched to the ground truth of retardance provided by the manufacturer (black line). Figure 2.c shows the orientation measurement before and after compensation. Before compensation, the measurement could not reveal different orientations set on the retarder at all. After compensation, the measured orientation matched the physical orientation of the retarder and the dynamic range was recovered to 180°. Figure 2.d shows the retardance weighted orientation image of a 4×4cm^2^ brain sample after birefringence compensation. The sample was index-matched to increase the signal-to-noise-ratio (SNR), which also increased the accuracy of orientation measurement^30^. The color-coded fiber orientation across this brain region was clear and matched with the imaging features revealing the fiber structures.

**Figure 2.**
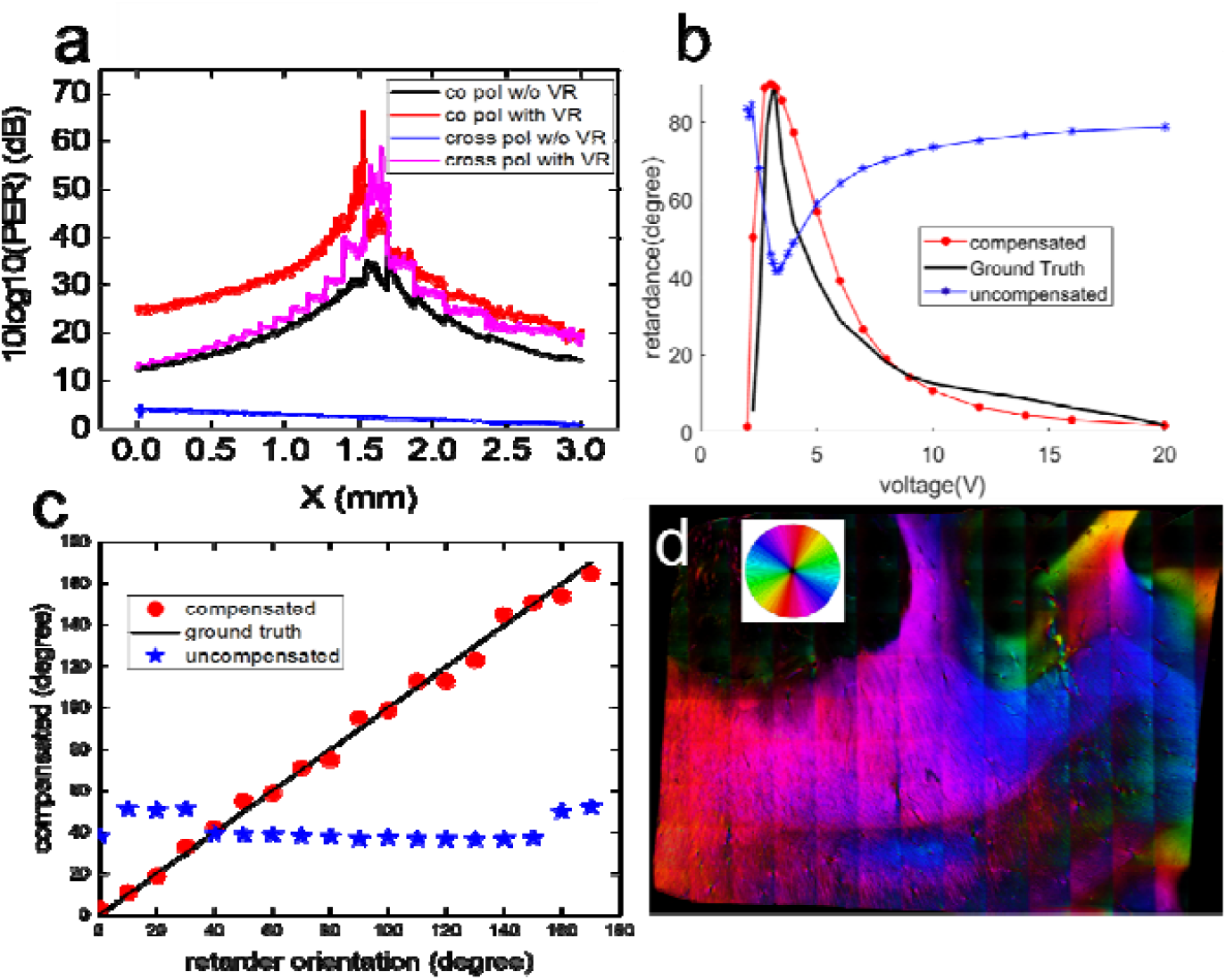
Characterization of polarization extinction ratio (PER), retardance and orientation before and after birefringence compensation using an electrically modulated variable retarder. (a) PER along the x axis at the center of FOV for cross and co polarization channels before and after compensation of birefringence. (b) Retardance measurement before (blue) and after (red) birefringence compensation using a variable retarder as a sample, compared with ground truth (black) of the retardance provided by manufacturer. (c) Comparison of orientation measurement of a retarder as a sample, before (blue) and after (red) birefringence compensation. Black line is the physical orientation of the retarder. (d) Retardance weighted fiber optic orientation map of a brain sample with index matching.

### 2.2 Myelin content

Optical properties estimated from PSOCT allow for quantification of myelin content. The scattering coefficient has been related to the myelin density across different human brain regions^31^. The back-scattering coefficient has been found to highlight the small fiber tracts and large neuromelanin pigmented neurons in the human midbrain^32^. The birefringence of myelin content has also been investigated and found to be related to myelin density and organization^10,33^. However, investigations connecting these mesoscopic optical properties of myelin are limited and validation at the microscopic scale has been missing. Here, in Figure 3.a-c we compared the estimated birefringence,, and for the same slice of the tissue block. Comparing these optical properties, we found all of them manifest a higher value in the white matter compared to the grey matter, owing to densely packed myelin content. All three contrasts revealed the cortical layers (yellow arrows in inset) in the grey matter, due to the variation of myelin density across the cortical areas. Comparing the line profiles across the cortical layers (Figure 3.d), we found that birefringence had the highest contrast separating those layers (indicated by the green dashed lines in d). We also found some differences among the three contrasts in the white matter, such as the sub-cortical U-fibers (indicated by the blue ROI and blue arrows) and some deep white matter regions (indicated by the red ROI). The microstructure of fibers underlying those optical property differences can be identified with high-resolution 2PM imaging. The U-fibers appeared to be brighter than the deeper fibers surrounding them in the birefringence and images. Zooming in to the higher resolution images of 2PM (Figure 3.e), we found the U-fibers were parallelly oriented axons (yellow arrows). This well-organized structure of the U-fibers could enhance the birefringence. In addition, the high wa concordant with previous studies^32^ showing that highlights small fiber tracts that were parallel to the imaging plane. For the red ROI in deep white matter, it was evident that the was higher than the surrounding area. Zooming into the corresponding area in 2PM, we found multiple large fiber tracts that obliquely ran through the imaging plane. Considering the strong scattering of densely packed fibers, the same structure may be the microscopic origin of the increased value. In short, our hybrid system allowed for the extraction of birefringence,, and from the same slice as well as the microscopic validation of the mesoscopic measurements using 2PM.

**Figure 3.**
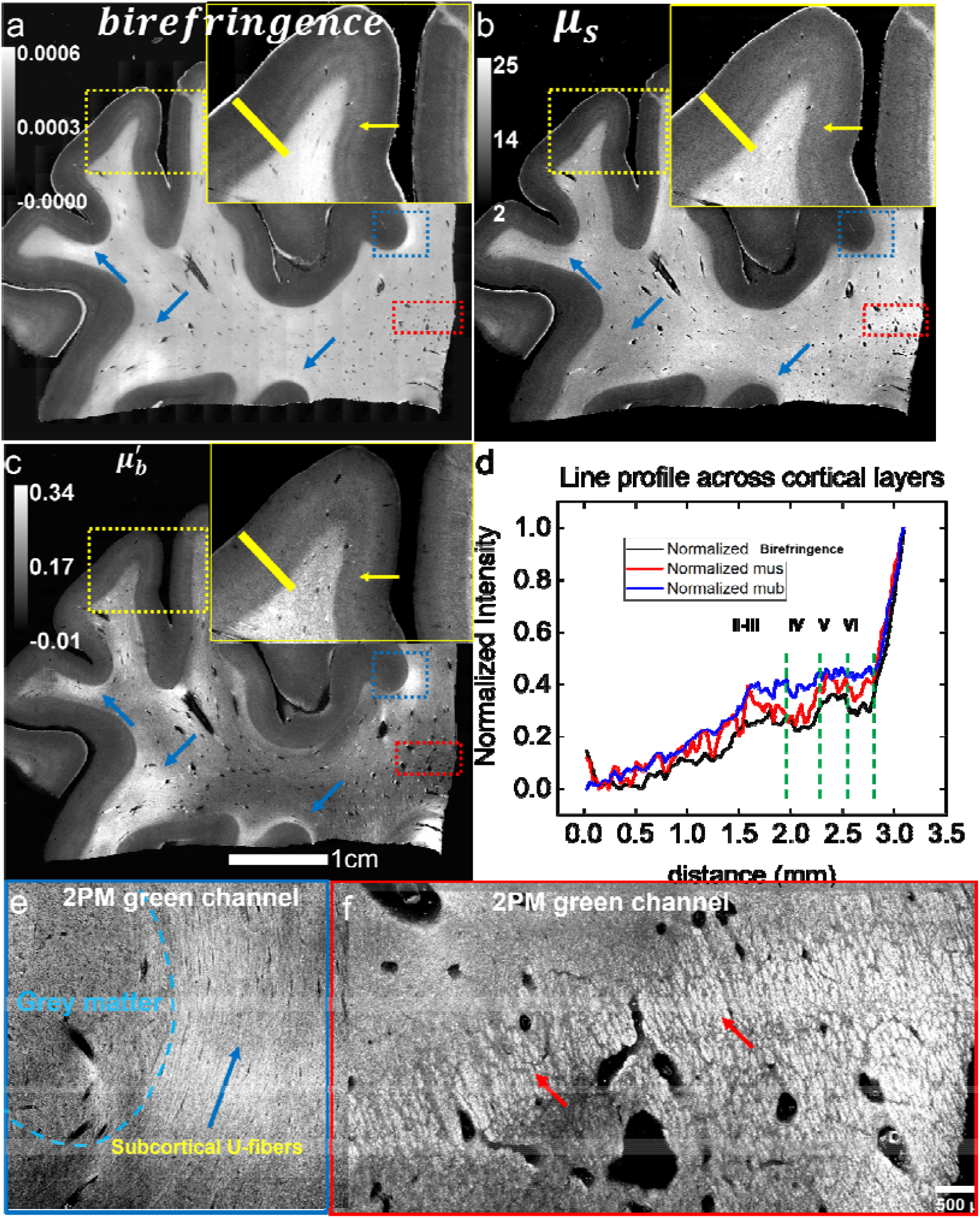
Comparison of different optical properties from PSOCT and the corresponding structure in 2PM. (a-c) Estimated birefringence (a), (b) and (c) from the same slice of a brain sample block. (d) normalized line profiles along the yellow bar in (a-c) inset that show the variation across cortical layers. Different cortical layers are separated by green dashed lines and denoted with layer numbers. (e-f) zoom-in of the blue and red ROIs by 2PM autofluorescence. (e) The white and grey matter boundary was indicated by the dashed lines. The blue arrows point to the parallelly oriented U-fibers underneath the cortex, which show high signals in the birefringence and 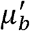 maps. (f) Red arrows point to the cross sections of densely packed myelinated fibers, corresponding to the increased scattering in the *μ*_*S*_ map.

### 2.3 Vascular structure

OCT has proven to be effective for imaging vascular structure of the human brain. Previous studies have shown quantitative analysis of the vascular geometry using graph theory following vessel segmentation^5,34^. However, due to the resolution limit and speckle noise, it has been challenging for OCT to visualize capillaries and small vessels smaller than 20 *μm* in diameter. Here, by combining 2PM, we enrichen the PSOCT vascular images with capillaries and details in the perivascular space. Figure 4.a shows a plane in the PSOCT volume where multiple vessels were captured, including large vessels in the white matter as the two yellow boxes highlight. Figure 4.b-e show the PSOCT and 2P image of the two ROIs in (a), where we found expansion of the perivascular space in vessels from both modalities (vessel walls indicated by red arrows). 2PM provided much higher resolution of the expanded perivascular space and a clearer image of the cross-section of vessel wall. Furthermore, we applied refractive index matching to another sample to reduce scattering and better visualize vessel structures. Figure 4.f shows a plane in the PSOCT volume from the sample that was index-matched and (g-j) show the zoom-in view of PSOCT and 2P images at the two yellow boxes. In (g) and (h) we found a twisting vessel squeezed into the perivascular space as well as two penetrating vessels from grey matter to white matter, as pointed out by the red arrows. Like the previous example, 2PM provided more details of the spiraling vessel and the branching of these penetrating vessels. The strong autofluorescence signal of these vessels might come from residue blood inside the vessel^22^. In (i), we found segments of vessels that don’t connect with each other, whereas in 2PM (j), we clearly found more vessels and capillaries creating a sophisticated network. We selected a small vessel (k) and measured the diameter to be around 15 *μm*, which would be challenging to visualize in the PSOCT image. In all, 2PM provided microscopic details that is complementary to the mesoscopic PSOCT vascular structure. What’s more, index matching reduced the scattering, and the residual hemoglobin boosted the autofluorescence signal of vessels, which strengthened our system’s ability to image details of the micro-vascular structure.

**Figure 4.**
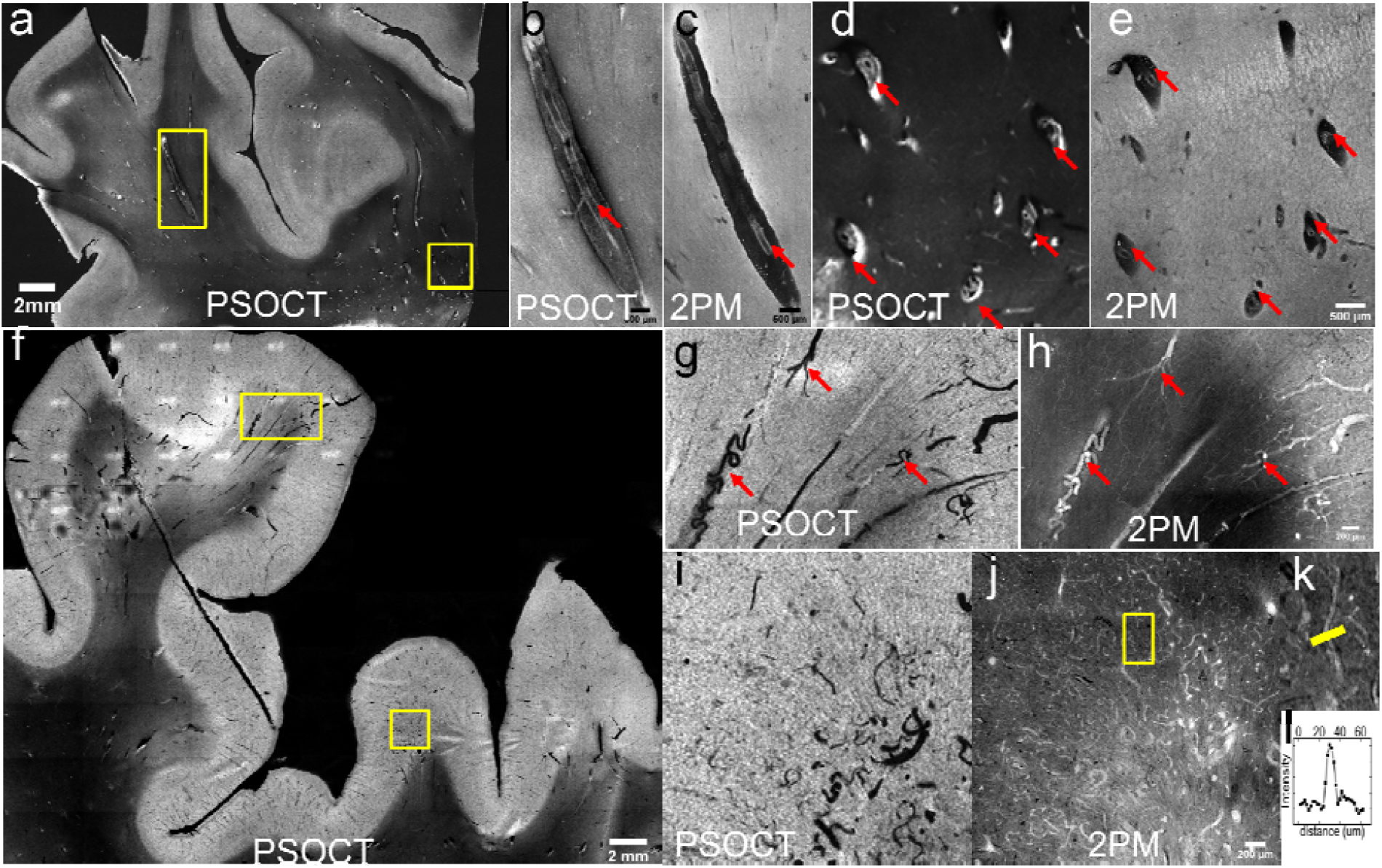
Vascular image from PSOCT and 2PM on two samples, one sample without refractive index matching (a-e) and the other sample with index matching (f-l). (a) A frame from PSOCT volume reconstruction that captures multiple segments of vessels. (b-c) Co-registered PSOCT and 2PM image of the vessel in the yellow box on the left of (a). The vessel wall is pointed out using red arrows. (d-e) PSOCT and 2PM images of the vessels in the yellow box on the right of (a). Cross-sections of five vessels are pointed out using red arrows. All these vessels in b-e demonstrate expansion of perivascular space. (f) A frame from PSOCT volume reconstruction of a sample after index matching. (g-h) PSOCT and 2PM images of the top box of (f). Two twisting vessel and a penetrating vessels from grey matter to white matter are pointed out using red arrows. (i-j) OCT and 2P images of the bottom box in (f). In 2PM image more small vessels and capillaries are visible. (k-l) zoom into the small vessel in yellow box in h and its cross-section line profile. The diameter of this vessel is approximately 15.

### 2.4 Lipofuscin and neurons

Lipofuscin has drawn increasing attention in the aging and degenerating human brain. The intraneuronal accumulation of lipofuscin is one of the most evident features in aged brain tissue^24^, where the distribution in the cerebral cortex^37,38^ and the role in neurodegeneration^39^ is intriguing for in-depth investigation. Our system utilized autofluorescence to provide a scalable mapping of lipofuscin across cortical layers. Figure 5.a shows the autofluorescence image from a 2×2cm^2^ brain sample using the long wavelength channel of our 2PM, which was dominated by lipofuscin autofluorescence. The grey matter was brighter than the white matter, which may be due to the higher density of lipofuscin in grey matter. Figure 5.b-c show the zoom-in of the left red box around the crown of the cortex and the segmented lipofuscin particles. We found densely distributed lipofuscin across the whole area (b), especially some larger lipofuscin particles as indicated by the red arrows. After segmentation, the large lipofuscin particles were easier to recognize. The lipofuscin in the superficial layer was sparsely distributed while in deeper layers small lipofuscin particles were densely distributed. In Figure 5.d-e, we show the lipofuscin distribution around the sulcus region in the right box of (a). Lipofuscin was densely scattered in all cortical layers (d). While examining the segmentation map (e), however, we also found layer structures enclosing different densities and sizes of lipofuscin within the grey matter, as indicated by the red arrows. It is noticeable that the background intensity of autofluorescence was inhomogeneous. Strong background autofluorescence reduced the contrast of lipofuscin and could cause error in the segmentation. For example, an empty hole (yellow arrow) was found in Figure 5.c.

**Figure 5.**
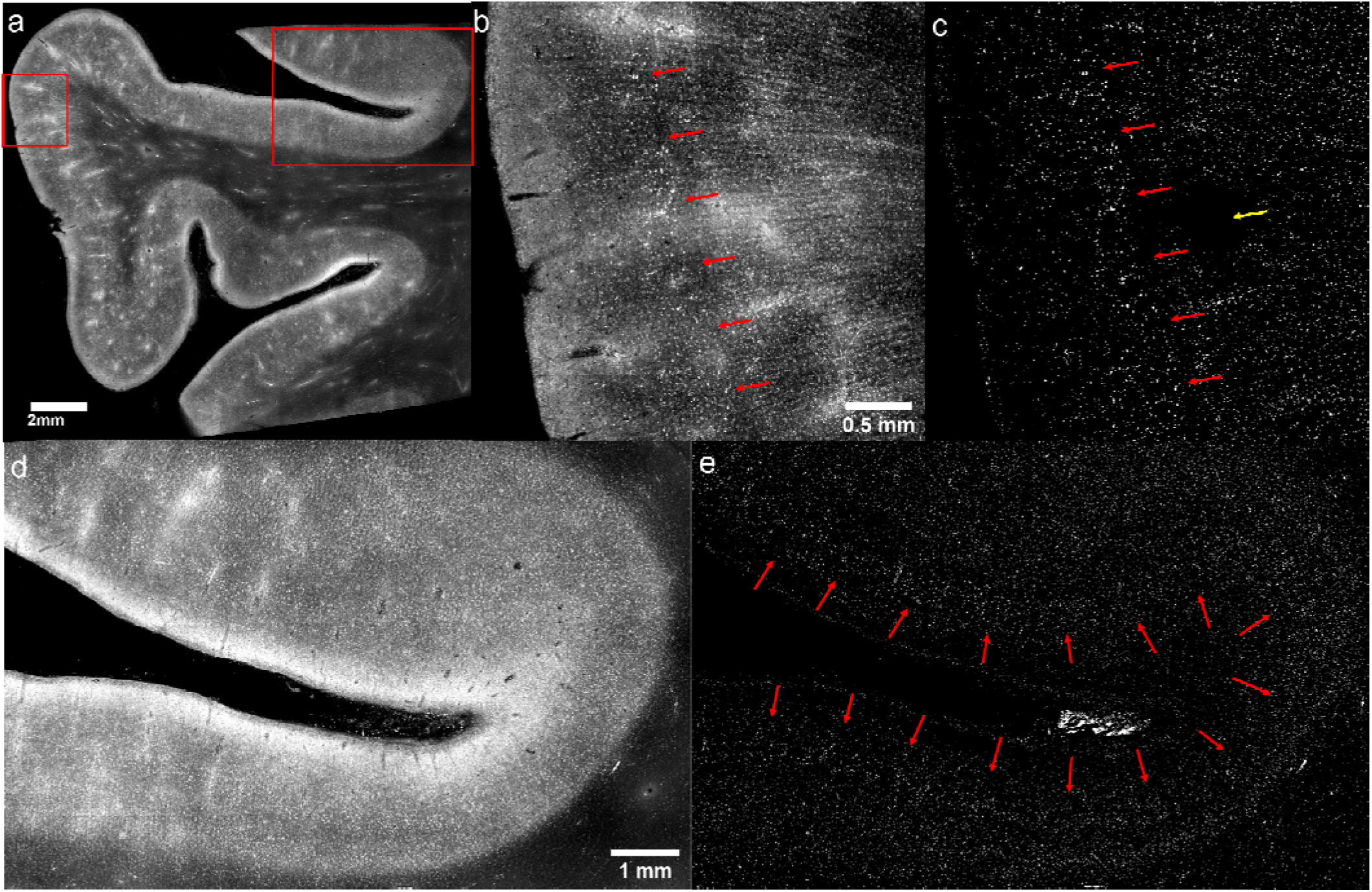
Autofluorescence image of lipofuscin from a brain sample and the segmentation of individual lipofuscin particles. (a) Overall intensity map of autofluorescence from a slice. (b) zoom into the left red box in (a) to show the lipofuscin distribution at the crown of cortex. Some larger lipofuscin particles form a layer in the grey matter and are pointed by the red arrows. (c) segmentation of the lipofuscin particles in (b). The larger lipofuscin particles were better visualized. An empty space is pointed out by the yellow arrow. (d) zoom into the right box in (a) to show the lipofuscin distribution around the sulcus of cortex. (e) Segmentation of the lipofuscin in (d). A dense lipofuscin layer was indicated by the red arrows.

We imaged another sample of cortex and examined the autofluorescence images from both the short and the long wavelength channels of our hybrid system. We found scattered lipofuscin in the long wavelength channel (Figure 6.a, red arrows) and dark spots in the short wavelength channel (Figure 6.b, yellow arrows). To better visualize the signals, we imaged the same region under a commercial 2PM (Bruker) at 0.5 resolution and found the same signals of lipofuscin and dark spots (Figure 6.c, yellow and red arrows). Benefiting from the high-resolution, we found that most lipofuscin were around the dark spots or the vessels. The shape of the dark spots appeared to be cell soma. To validate, we sectioned a 30 slice after imaging with a Bruker 2PM and performed a Nissl stain on the same slice, which was a standard method for visualizing cell soma. We selected a region enclosing layers II, III and IV of cortex where a high density of neurons was expected, and investigated neurons in three ROIs (Figure 6, second row: Nissl; third row: Bruker 2PM). We found the markers of lipofuscin and dark spots in the 2PM images agreed with the neurons in the Nissl stain at the same location (red and white arrows). Therefore, 2PM autofluorescence combining lipofuscin and dark spots can be used to quantitatively identify the neurons. Two observers counted the cell number in the Nissl stain and in the autofluorescence image (Figure 6, bottom row). Both observers found similar counts of neurons in the Nissl and 2P images, and the correspondence rate between the two modalities was 92% on average.

**Figure 6.**
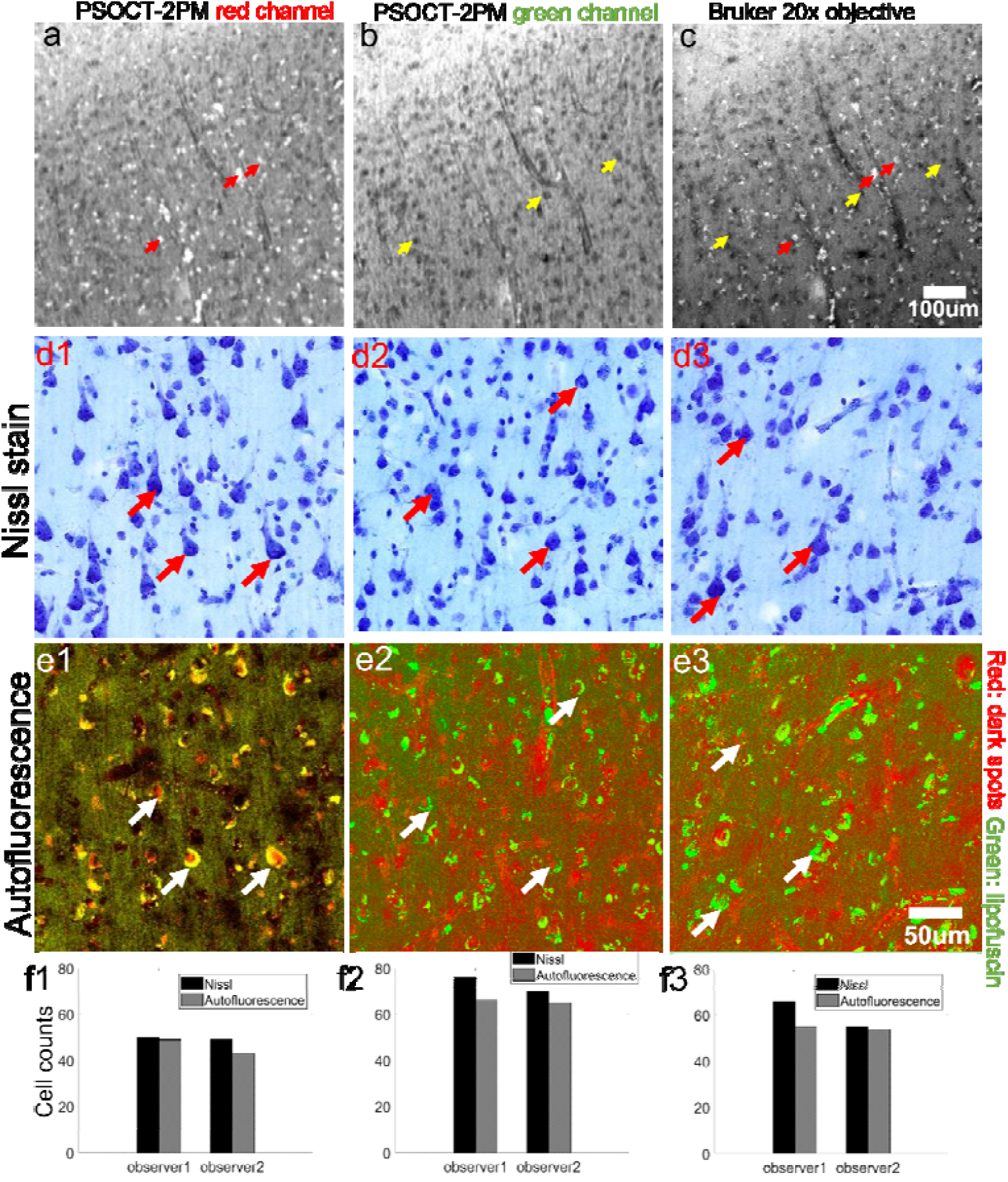
Autofluorescence images of brain tissue at different resolutions, and comparison with Nissl stain. (a) Long wavelength channel of the PSOCT-2PM system showing the lipofuscin autofluorescence. (b) Short wavelength channel of the PSOCT-2PM system showing the dark spots. (c) 0.5 high resolution image obtained with a 20x objective from the Bruker 2PM system. (d1-d3) Three ROIs in the Nissl stain image. ROIs were taken from the grey matter. (e1-e3) Bruker 2PM images of corresponding ROIs as in d1-d3, with lipofuscin in the green channel and inverted dark spots in the red channel. A few representative neurons in the Nissl stain and the corresponding dark-spot-lipofuscin structures are highlighted (red and white arrows). (f1-f3) Manual counts of neuron numbers from the two modalities by two observers.

## 3. Discussion

We have developed a hybrid system that combines serial sectioning PSOCT and 2PM and generates simultaneous multi-contrast imaging of large volumes of human brain tissue. The 3×3mm^2^ large FOV of both PSOCT and 2PM reduced the imaging down-time during stage translation between tiles and improved the throughput. We intentionally designed the 2PM to match the acquisition speed of PSOCT, while allowing a microscopic investigation of the sample. The balanced resolution of the integrated system allowed for visualization of multiple fine structures, all using intrinsic optical properties. Additionally, because the two microscopes imaged the same sample simultaneously with largely overlapped FOV, the need for image registration in post-processing was markedly reduced. Although previous studies have combined OCT with 2PM to enrich the image contrasts of the individual modalities^18,24,25^, one of the challenges of combining PSOCT with 2PM is system birefringence that prevents accurate measurement of sample retardance and orientation. System birefringence originates from the optical telescopes, dichroic mirrors, and scanning mirrors, and results in a substantial bias to the polarization measurements of the sample. We removed this bias by using an adjustable retarder that enabled measuring true retardance and orientation from the sample. As a result, we were able to investigate multiple signals of scattering, birefringence, and autofluorescence from the human brain at different scales. Utilizing these contrasts, we enrichened the measurements of the myelin content, vascular structure, lipofuscin, and neurons in volumetric brain tissue blocks.

For myelin content, we estimated three optical parameters from PSOCT for investigation of myelin density and used high-resolution 2PM as microscopic corroboration. We found that both birefringence and 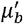 highlighted the subcortical U-fibers (Figure 3.a-c), while *μ_s_* was sensitive to myelin density in deep white matter. We used 2PM images to corroborate the above measurements. The highly organized U-fibers and densely packed fiber bundles of the deep white matter were recognized by the high resolution autofluorescence of 2PM. The different organizations of fibers in these areas might contribute to variation in these optical parameters. We demonstrated that by comparing the mesoscopic measurements from PSOCT with the microscopic information from 2PM, our system reliably measured human brain myeloarchitecture. Additionally, our system removed birefringence bias from the optical components of the system, generating accurate optic axis orientation maps, which were comparable to standalone PSOCT^10^.

For vascular structure, PSOCT and 2PM provided complementary information at different resolutions, that enrichened the vascular images obtained from the individual modalities. Specifically, PSOCT reconstructed 3D vascular structures in a large volume, allowing for vessel tracking and segmentation at the mesoscopic resolution. Yang et al^6^ segmented vascular structures from the OCT volumetric images and developed quantitative metrics for characterizing the geometric properties of vascular structures. 2PM provided microscopic images that revealed details about the expanded perivascular space, twisting vessels, penetrating vessels, and the intricate capillary network (Figure 4). Spiraling vessels^40,41^ and widening of perivascular spaces are important aspects of vascular morphology that contribute to small vessel disease^42^, aging^43^ and Alzheimer’s disease^44,45^. Our hybrid system added autofluorescence as an additional contrast and enabled the recognition of vascular abnormalities at resolutions that were beyond that of OCT alone.

Lipofuscin, a pigmented substance found within lysosomes or the cytosol of aging cells, has attracted increased attention in the scientific literature as a feature of degeneration. We demonstrated the visualization of lipofuscin using autofluorescence in the 500-700nm wavelength channel and developed a segmentation method to segment lipofuscin particles from background. The segmented lipofuscin maps could be used for future investigation of lipofuscin density and size distribution across cortical layers^38^ and can be applied to large scale volumetric brain images from different regions of the human brain. This method may be useful to investigate differences in lipofuscin distribution between different neurodegenerative processes, as previous investigations were limited to small sample sizes of only one hundred of neurons^46,47^.

Lipofuscin was not the sole source of autofluorescence; elastin and collagen in the extracellular matrix^29^ and formalin fixation^48^ also contribute to autofluorescence. Interestingly, we found dark spots in the 435-485nm wavelength channel (Figure 6) that colocalize with lipofuscin. By comparing with Nissl stain we found this lipofuscin-dark-spot structure shows strong correspondence with neuron soma, suggesting that the method might be useful as a neuron counting strategy ^49^. Although the validation and quantification of autofluorescence was conducted at sub-micro resolution by a separate 2PM in this study, recent advancements in machine learning methods show promise as a means of segmenting low-resolution images using prior knowledge from high-resolution ones^50,51^.

There are several limitations to our method. First, while combining 2PM with PSOCT allowed for the visualization of smaller vessels and capillaries, 2PM was performed at a single depth integrating over 48 *μm* of the tissue, about one third of PSOCT depth of focus. Therefore, there might be information mismatch between the co-registered PSOCT and 2PM images. This imaging strategy also precluded a full volumetric reconstruction from 2PM images. To overcome the problem, depth scanning is needed at a cost of longer acquisition time. Second, to resolve even smaller structures such as individual axons, myelin sheaths, the full capillary bed, and small cells, a sub-micron resolution is necessary.

In summary, our method enabled scalable and multi-resolution investigation of blood vessel structures, myelin content, lipofuscin, and neuron density in the human brain. In addition, our method allowed their simultaneous spatial correlation in the human brain. Previous studies have observed neuron loss, myelin degradation, and vascular changes such as spiraling vessels and altered small vessel density in the context of cerebrovascular disease, Alzheimer’s disease, and aging^44,52–57^. However, the relationships among neuron loss, myelin degradation, and vascular abnormalities in the human brain and their roles in aging and disease is still unclear. Our method, which provides volumetric and distortion-free reconstruction of large volumes of human brain tissue offers a powerful tool to further understand neurological disorders and neurodegeneration.

## 4. Methods

### 4.1 Sample

Five human brain samples were used in this study. Four neurologically normal samples were obtained from the Massachusetts General Hospital Autopsy Suite, and one with Stage VI Alzheimer’s disease was obtained from Boston University Alzheimer’s Disease Research Center brain bank. These samples were fixed by immersion in 10% formalin for at least two months ^51^. The post-mortem interval did not exceed 24 h. The samples were washed in phosphate buffered saline for a month to remove residual fixation agents and then embedded in 4.5% agarose for tissue support^52^. During embedding, the brain blocks were warmed to 65 °C to allow complete penetration of agarose into the deep sulcus. A 3D printed base plate was embedded onto the bottom of the agarose block and screwed to the bath container to mount the tissue block during the one-week serial block-face imaging (inset in Figure 1). Two samples were refractive-index matched in 60% 2,29-thiodiethanol (TDE)^58^. The index-matching procedure was described in detail previously^58,59^. The other samples were immersed in DI water during imaging. The immersion liquid was changed every day to remove the debris from cutting that could degrade the image quality.

### 4.2 PSOCT-2PM system

The hybrid system consists of the PSOCT, 2PM, a custom-built vibratome, XYZ motorized stages, and the control software for the whole system. The PSOCT microscope uses a swept light source (AxsunTech) with 100kHz swept rate, a central wavelength of 1310nm, and a spectral full width half maximum (FWHM) of 110nm, yielding an axial resolution of 5.6 *μm* in brain tissue (n=1.4). At the interferometer, a polarization controller (Thorlabs Inc) and a polarization beam splitter (PBS) ensure maximum power input with horizontal polarization. A 50:50 beam splitter (BS) separates the light intensity equally onto the sample arm and reference arm. In the sample arm, an achromat quarter-wave plate (QWP) is placed in front of the sample arm at 45 degrees to generate circular polarization on the sample. The scanning path consists of a pair of galvo mirrors (Thorlabs Inc), a 4x telescope and a 4x objective (Olympus, UPLFLN4x, NA=0.13), providing 5 *μm* lateral resolution and a 150 *μm* confocal parameter. In the reference arm, a linear polarizer (LP) is placed in front of the reference arm at 45 degrees to make the light intensity equal between horizontal and vertical polarization states. A retroreflector (RF) is placed at the end of the reference arm to back-reflect the light. The back-reflected light from the sample arm and the reference arm interferes when re-combined in the interferometer and is split by two PBS into horizontal and vertical polarization states, which, through balanced detection, result in the co- and cross-polarization measurement of the interference pattern. To acquire the spectrum in even k-space, the k-clock of the light source is fed into a high-speed digitizer (ATS9350, AlazarTech) as a sampling clock. A Graphic Processing Unit (RTX4000, NVIDIA) is used to perform real-time FFT, and the spatial-domain data is trimmed to only save the first 1mm depth. The post-objective power was measured to be 3.7mW, achieving a 95dB SNR in both polarization channels. A 3×3mm^2^ FOV was used with 3 *μm* lateral step size and 10% overlap between tiles.

The optical components in the sample arm generated a large system birefringence that caused substantial errors in the retardance and optic axis orientation measurements. To address this problem, we used a variable retarder (VR) in the sample arm, assuring the measurement to be pure tissue birefringence (Figure 2). The compensation process can be carried out in 3 steps:

1. We imaged a silver mirror and calculated the retardance. Assuming there was no system birefringence, the measured retardance should be 0. So, the none-zero retardance we measured was the accumulated retardance from the optics.
2. We set the retardance of the variable retarder to be opposite of the value we calculated in step 1, placed it before the galvo mirrors and rotated it to minimize the signal from mirror in the cross-polarization channel.
3. We then fine-tuned the retardance and orientation of the variable retarder iteratively to minimize the cross-polarization channel signal of sliver mirror.

The 2PM used a Ti:Sapphire mode-locked laser with 80MHz rep rate, 100fs pulse duration and approximately 3W laser power at 820nm. Right after the laser output, an attenuator made of a motorized half-wave plate (HWP) and a PBS was used to modulate the power input to the microscope system. Actual power on the sample was 128mW. Following the attenuator, a 1.25x beam expander combined with the 4X telescope in the scanning path expand the beam size to fill the back-aperture of the objective. The 2PM beam and PSOCT beam in the sample arm were combined using a dichroic mirror above the objective. Sharing the same objective with PSOCT, 2PM achieved a 2 *μm* lateral resolution and 48 *μm* axial resolution. The FOV and the imaging speed of 2PM were matched to those of PSOCT, as a result, we used a 2 *μm* lateral step size. The depth of focus of PSOCT and 2PM were carefully adjusted to overlap, assuring that they imaged the same region of the brain tissue. However, because 2PM had shorter depth of focus, 2PM only integrated approximately one third of the depth imaged by PSOCT. Two detection channels were available, the short wavelength channel covered from 435 to 485nm and the long wavelength channel covered from 500 to 700nm to capture lipofuscin autofluorescence.

Two high-performance computers controlled the PSOCT and 2P system separately, which were synchronized using frame start and stop triggers. For the 2PM system, we used the ScanImage (Vidirio Inc) to control the 2PM acquisition. For the serial sectioning PSOCT system, we wrote custom software in LabVIEW to automate the PSOCT acquisition, XYZ stages and vibratome slicing. PSOCT system generated tens of TBs volumetric data, leading to the demand for large storage space and high computing power to process the data parallelly. We established a 10Gb/s Ethernet connection between the PSOCT acquisition computer, a local server with 70TB RAID10 disk space, and the remote Boston University Shared Computing Cluster (BU SCC), with the local server playing as a transfer station that separated the acquisition computer from data saving and transfer. Post-processing was performed on SCC clusters.

### 4.3 System characterization

We characterized the lateral and axial resolution of both microscopes (Supplementary Figure 1). Using 1 *μm* fluorescent beads, the lateral and axial resolution of 2PM was measured to be 2 *μm* and 48 *μm*, respectively (a, b). The lateral resolution of PSOCT was 5 *μm*, where the modulation transfer function showed 50% contrast between the line-pair (c). Using the air-glass interface and considering the brain tissue index of refraction (n=1.4), the axial resolution of PSOCT was measured 5 (d). The confocal parameter was measured to be 150 *μm* using an intralipid phantom (Supplementary Figure 2).

In swept source OCT, the sensitivity drops along depth due to the limited spectral resolution of the swept light source, yielding a sensitivity roll-off. Supplementary Figure 1.e shows the sensitivity roll-off measurement at different reference mirror positions. We found the sensitivity remained unchanged over the first 1 mm depth. The purple horizontal line represented the fitting of the peak intensities using equation 2 in ^60^. As the brain signals were captured within the first 1 mm, the small sensitivity roll-off relieved us from performing roll-off compensation during post-processing.

We performed a thorough characterization of the polarization measurements in PSOCT before and after compensating the system birefringence. To obtain the polarization extinction ratio (PER, Figure *2*.a), we used a microscope glass slide as sample and rotate the QWP in the sample arm to get maximum and minimal signals from the cross and co polarization channels. PER was then calculated by ratio of the maximum and the minimum intensity signal from the same channel. To characterize the retardance and orientation measurements (Figure *2*.b-c), we used a second variable retarder as the sample and imaged the bottom surface of the retarder. We adjusted the voltage of the retarder to change the retardance and rotated the retarder to change the optic axis. We then compared the PSOCT-measured retardance and orientation with the true retardance provided by the manufacturer and the actual orientation of the retarder.

### 4.4 Image acquisition and post-processing

#### 4.4.1 Data acquisition

Block-face imaging and serial sectioning were performed on three human brain blocks. For the sample without index matching, PSOCT imaged 150 *μm* deep into the tissue with 5 *μm* resolution, while 2PM acquired a single plane integrating a depth range of 48 *μm* (axial resolution). Then a 150 *μm* thick slice was cut off by the vibratome. During serial sectioning, we also cut 30 *μm* thick slices for Nissl stain. For the sample with index matching, PSOCT could effectively image 450 *μm* deep into the tissue. We achieved this by imaging the sample at two different focal depths, separated by 225 *μm*. Two 2PM images were taken at the same time. Then a 450 *μm* thick slice was cut by the vibratome.

#### 4.4.2 Distortion correction

The optical distortions can be categorized into three categories. The first two are grid distortion and field curvature distortion, which are geometric distortions along the lateral and axial directions, respectively. Grid distortion appears as a warped image along the lateral dimensions due to the non-telecentricity of the optical system and is present in both PSOCT and 2PM images. Field curvature distortion appears as a curved depth profile in the OCT B-line image due to the optical path length difference at different scan angles. The third category of distortion is shading distortion, which appears as non-uniform signal intensity across the FOV due to the vignetting of light beams along the optical system. Shading distortion is present in both modalities. Since the two geometric distortions depend purely on the optical system, pre-calibration can be applied to correct them. Briefly, a grid target (Thorlabs Inc) parallel to the table surface was imaged before the experiment. The back-reflection signal off the top surface shared the same field curvature as any well-cut sample. A 2D look-up table of the curvature was obtained from the grid target and applied to the brain images to flatten the signal surface. Next, the average intensity projection (AIP) of C-scan images of the grid target along depth provided the warped image of the grid that was used in grid distortion correction. An ideal grid pattern was generated in MATLAB with the same pixel spacing and orientation as the captured image. Using the UnwarpJ plugin of ImageJ, the warped grid pattern was registered to the non-distorted pattern and a transformation matrix was generated to unwarp the sample images. A similar procedure was applied to correct the 2PM images. Note that for 2PM, a uniform fluorescent glass slide was placed under the grid pattern slide, with the side coated with the grid facing downwards to get a fluorescent grid image.

For the shading distortion, we employed the BaSiC shading correction algorithm^61^, using the ImageJ plug-in.

#### 4.4.3 Stitching and volume reconstruction

The hybrid system generated two sets of data, the speckle-noise contaminated PSOCT images and the autofluorescence images. As the two modalities were acquired synchronously, the same coordinates could be used to stitch both images. Because 2PM images had better SNR, we stitched 2PM images to get stitching coordinates and then use them to stitch PSOCT images. We used the stitching plugin of ImageJ to find the optimal stitching coordinates for the 2PM images. We selected three planes separated at different depths of the volume, merging them into a RGB format and ran the stitching plugin to find the global optimal coordinates for stitching and applied them for all the slices, which were then stacked to reconstruct the volume.

#### 4.4.4 Scattering and birefringence fitting

Extraction of scattering coefficient^32,59^, back-scattering coefficient and birefringence^30^ followed previously reported methods, by fitting the depth profiles of PSOCT intensity and retardance.

#### 4.4.5 Registration between PSOCT and 2PM images

As PSOCT and 2PM were acquired simultaneously, and all distortions have been corrected in previous steps, registering these two modalities only need to scale PSOCT and 2PM to the same pixel size and then find out how many pixels were shifted between the two FOVs in X and Y dimensions. We determined this shift by finding the maximum overlap between the two images on the same vessel. The scaling factor from PSOCT to 2PM was 1.5. The shift from PSOCT to 2PM was -50 pixels in X dimension and 113 pixels in Y dimension.

#### 4.4.6 Histology process and imaging

The Nissl staining protocol was described previously^49^. We used a microscope slice scanner (Olympus, VS200) for imaging the Nissl-stained slices. We used a 20x objective, with a lateral resolution of 0.5 *μm*. The commercial software VS200 performed intensity correction and stitching automatically.

#### 4.4.7 Bruker 2PM imaging

After imaging under our hybrid system, we took the sample to a Bruker 2PM for high resolution imaging. We used a 20x water objective, yielding a lateral resolution of 0.5 *μm* and axial resolution of 1.3 *μm*. We used 820nm excitation wavelength, the same as in the hybrid system. The emission filter was 500-550nm, revealing both the lipofuscin and the dark spots as shown in the hybrid system. Z-stacking was carried out in 10 *μm* steps and a total of 30 *μm* depth was imaged. Two-dimentional autofluorescence images were generated from the Z-stack. Lipofuscin map was obtained by the maximal intensity projection of Z-stack and the dark spot map was obtained by the minimal intensity project.

### 4.5 Segmentation of lipofuscin

Segmentation of lipofuscin was based on high-pass filtering of the image and adaptive thresholding afterwards. Specifically, the process was carried out in the following steps:

a. Low-pass filtering was applied on the 2PM long wavelength channel using the Gaussian smoothing function of ImageJ. We used Sigma (radius) value of 50, which corresponded to 100 *μm*.
b. The original image on the 2PM long wavelength channel was divided by the filtered image in (a). This way we removed the low frequency background.
c. The image was binarized using the Threshold function of ImageJ and Huang method on the image in (b). We used the lower bound 1.4, which gave best discrimination between lipofuscin and background.

## Acknowledgment

We thank Ms. Ana Vitantonio and Prof. Douglas Rosene in the Boston University Medical campus for the help with Nissl staining. We thank Prof. Martin Thunemann from Boston University for the guidance in immunohistochemistry. We thank the Micro and Nano Imaging (MNI) core facility of Boston University for the imaging of Nissl and DAPI (not shown) stained slices. We thank Mr. James Goebel from the IT department of Boston University for setting up the network and disk space of the server for extensive data transfer and storage. We thank Ms. Mingshan Guo for the manual counting of neurons in Figure 4.

Support for this research was provided in part by the BRAIN Initiative Cell Census Network grant U01MH117023, the National Institute for Biomedical Imaging and Bioengineering (P41EB015896, 1R01EB023281, R01EB006758, R21EB018907, R01EB019956, P41EB030006, R00EB023993), the National Institute on Aging (1R56AG064027, 1R01AG064027, 5R01AG008122, R01AG016495, 1R01AG070988), the National Institute of Mental Health (R01 MH123195, R01 MH121885, 1RF1MH123195), the National Institute for Neurological Disorders and Stroke (R01NS0525851, R21NS072652, R01NS070963, R01NS083534, 5U01NS086625,5U24NS10059103, R01NS105820, R01NS128843), a NIH career development award R00EB023993 to HW and grant number 2019-189101 from the Chan Zuckerberg Initiative DAF for CM. Additional support was provided by the NIH Blueprint for Neuroscience Research (5U01-MH093765), part of the multi-institutional Human Connectome Project. And the Boston University National Science Foundation Research Traineeship Program (NRT) Understanding the Brain: Neurophotonics (NSF NRT UtB: Neurophotonics) and was made possible by the resources provided by Shared Instrumentation Grants 1S10RR023401, 1S10RR019307, and 1S10RR023043. Fundings associated with ACM include U54 NS115266/NS/NINDS NIH HHS/United States, P30 AG072978/AG/NIA NIH HHS/United States, U19 AG068753/AG/NIA NIH HHS/United States, P30 AG072978/AG/NIA NIH HHS/United States. In addition, BF has a financial interest in CorticoMetrics, a company whose medical pursuits focus on brain imaging and measurement technologies. BF’s interests were reviewed and are managed by Massachusetts General Hospital and Partners HealthCare in accordance with their conflict of interest policies.

## Supplementary Figures

**Supplementary Figure 1.**
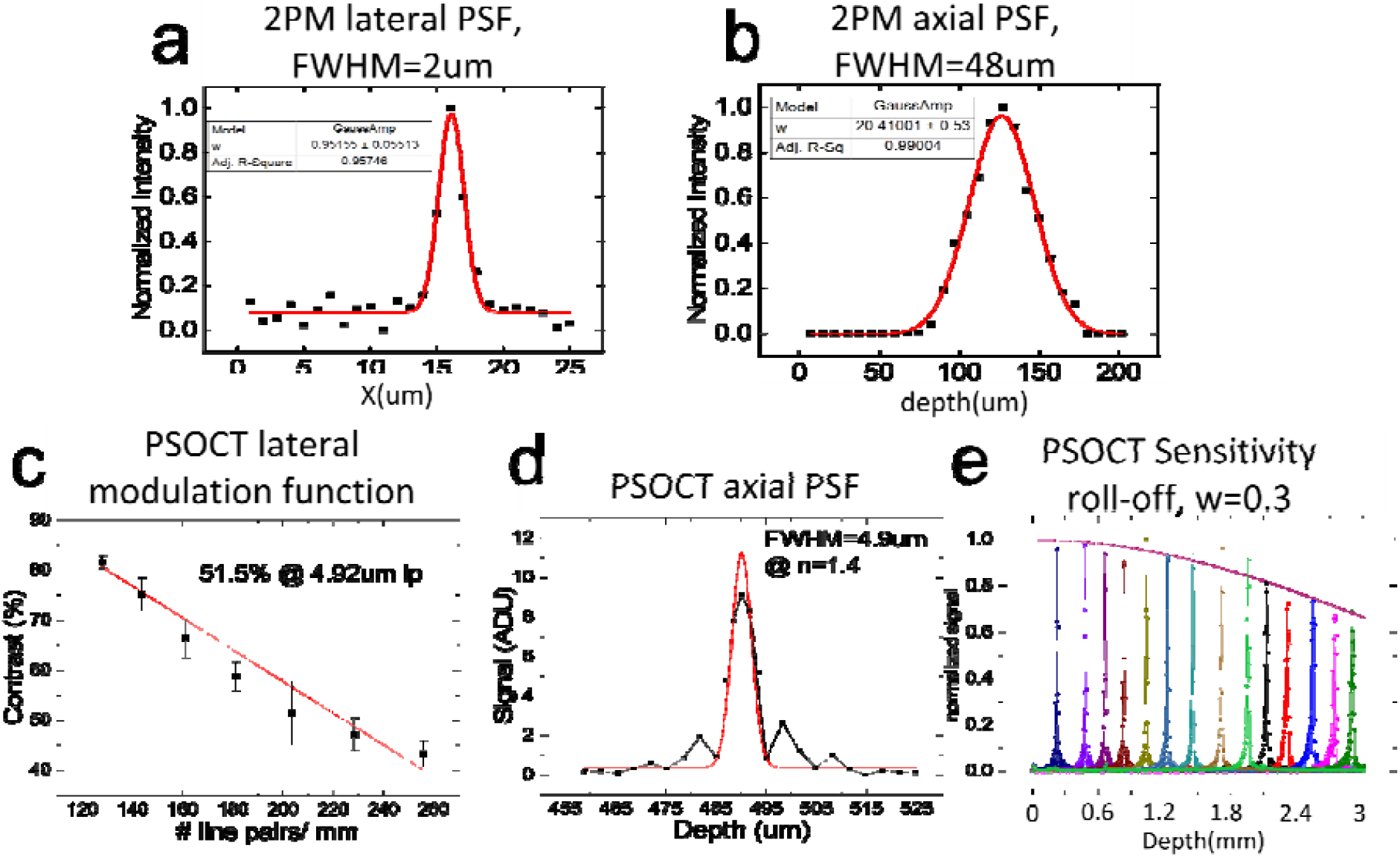
Lateral and axial resolution measurement of the PSOCT-2PM system. (a) lateral resolution of 2PM measured to be 2 using 1 diameter fluorescent beads. (b) axial PSF of 2PM measured to be 48. (c) Lateral modulation function of PSOCT system using airforce target shows 50% contrast at 5 line pair. Red line shows linear approximation of theoretical values. (d) Axial PSF of PSOCT measured to be 5 using glass slide surface. (e) Sensitivity roll-off of the PSOCT signal over depth.

**Supplementary Figure 2.**
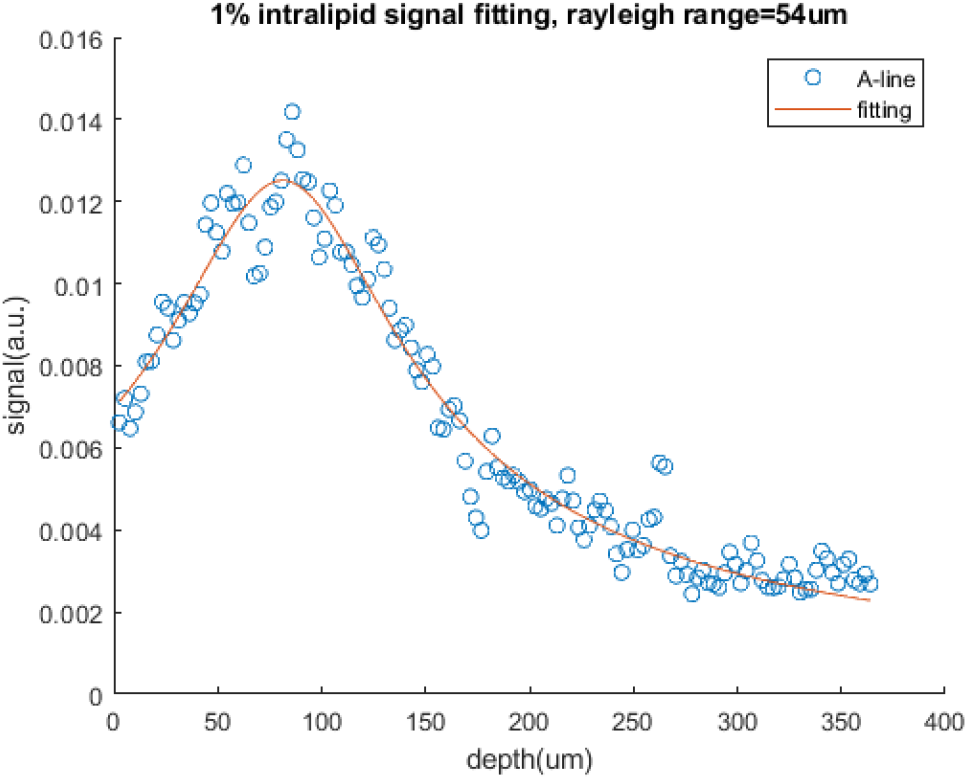
Four parameter optical fitting of 1% volume concentration intralipid solution. Rayleigh range was estimated to be 54 *μm*, which corresponds to 108 *μm* of confocal parameter. With higher scattering, effective Rayleigh range would increase^59^, we found in brain tissue the confocal parameter is about 150 *μm*.

